# Evaluating the Performance of Widely Used Phylogenetic Models for Gene Expression Evolution

**DOI:** 10.1101/2023.02.09.527893

**Authors:** Jose Rafael Dimayacyac, Shanyun Wu, Daohan Jiang, Matt Pennell

## Abstract

Phylogenetic comparative methods are increasingly used to test hypotheses about the evolutionary processes that drive divergence in gene expression among species. However, it is unknown whether the distributional assumptions of phylogenetic models designed for quantitative phenotypic traits are realistic for expression data and importantly, the reliability of conclusions of phylogenetic comparative studies of gene expression may depend on whether the data is well-described by the chosen model. To evaluate this, we first fit several phylogenetic models of trait evolution to 8 previously published comparative expression datasets, comprising a total of 54,774 genes with 145,927 unique gene-tissue combinations. Using a previously developed approach, we then assessed how well the best model of the set described the data in an absolute (not just relative) sense. First, we find that Ornstein-Uhlenbeck models, in which expression values are constrained around an optimum, were the preferred model for 66% of gene-tissue combinations. Second, we find that for 61% of gene-tissue combinations, the best fit model of the set was found to perform well; the rest were found to be performing poorly by at least one of the test statistics we examined. Third, we find that when simple models do not perform well, this appears to be typically a consequence of failing to fully account for heterogeneity in the rate of the evolution. We advocate that assessment of model performance should become a routine component of phylogenetic comparative expression studies; doing so can improve the reliability of inferences and inspire the development of novel models.

## Introduction

While DNA holds the genetic information required for life to work, other elements are largely required for cells to function. These functional elements are responsible for the molecular processes that eventually lead to phenotypes [Kellis et al., 2014]. The most prominently studied of these elements is gene expression. There is a long tradition of thinking about gene expression evolution in a comparative context [Gilad et al., 2006, King and Wilson, 1975, Wray, 2007], yet it is only recently that it has been feasible to gather gene expression data for multiple species in a standardized way. This has opened up new avenues for investigating the evolutionary processes responsible for generating diversity [Hill et al., 2021, Price et al., 2022, Smith et al., 2020] of changes in gene expression. Identifying interspecies differences in gene expression can pinpoint which sets of genes are responsible for differences between organisms. Many such studies have used the approach of directly comparing gene expression levels between orthologs to understand an array of topics, such as the function of epigenetic modifications [Cain et al., 2011], the connection between DNA and methylation [Hernando-Herraez et al., 2015], and the evolution of enhancer regions [Villar et al., 2015]. The studies mentioned above (in addition to many others in the field) use pairwise comparisons in which all gene expression values from all species are compared to one another. Essentially, this assumes that gene expression values from different species all represent independent measurements [Dunn et al., 2018]. However, due to their shared evolutionary history, more closely related species will resemble each other in many ways and some of these shared (and, in many cases, unmeasured) attributes will influence how focal variables (here, gene expression and some attribute of interest) are associated with one another [Felsenstein, 1985, Uyeda et al., 2018]. While this challenge has been widely recognized across the biological sciences, many comparative gene expression studies still do multi-species comparisons with sequential pairwise comparisons, which a recent study demonstrated could be highly misleading [Dunn et al., 2018].

In addition to controlling for unobserved (and phylogenetically structured) confounding variables, phylogenetic comparative methods (PCMs; for reviews of these methods see Pennell and Harmon [2013] and Harmon [2019]) are increasingly being used to characterize the evolutionary dynamics of gene expression over time, for example, by looking for the signature of selection in the distribution of gene expression values at the tips [Bedford and Hartl, 2009, Brawand et al., 2011, Dunn et al., 2013, Oakley et al., 2005, Price et al., 2022, Rohlfs et al., 2014, Rohlfs and Nielsen, 2015]. And accordingly, there have been a number of recent methodological developments, including computational platforms for simulating [Bastide et al., 2023] and analyzing [Bertram et al., 2023] phylogenetic comparative gene expression datasets.

While this work is tremendously exciting, it is important to note that the reliability of the inferences from phylogenetic comparative methods hinge upon the performance of the phylogenetic model that is fit to the data [Boettiger et al., 2012, Brown and Thomson, 2018, Garland et al., 1992, Pennell et al., 2015, Price, 1997, Uyeda et al., 2021]. There is a long tradition of using PCMs for modeling the evolution of morphological and ecological phenotypes but as comparative, multi-species gene expression datasets are starting to become more available, the performance of the models in this new context is not well understood. And there are reasons to think that results from applying phylogenetic models to well-studied morphological phenotypes might not apply to gene expression data. First, evolutionary models of continuous traits were derived under the assumptions of quantitative genetics, where phenotypes are controlled by a large (effectively infinite) number of loci [Felsenstein, 1988, Hansen and Martins, 1996, Lande, 1976, Lynch, 1990, Pennell and Harmon, 2013, Turelli, 1988]. We might expect the expression level of a given gene to behave less like an idealized polygenic trait owing to the outsized importance of the *cis*-regulatory region in determining the expression level [Dhar et al., 2021, Fuso et al., 2020, Matharu and Ahituv, 2020, Romero et al., 2012]. On the other hand, searches for eQTLs have turned up a moderately large number of candidate loci potentially involved in the regulation of some genes [GTEx Consortium, 2020, Hill et al., 2021, Rockman and Kruglyak, 2006]. Theoretical work has demonstrated that differences in the genetic architecture of traits influence the distribution of phenotypes among species [Schraiber and Landis, 2015]. Second, unlike traits such as height or mass, where the meaning of a measurement is straightforward, this is not the case for gene expression [Diaz et al., 2023]; the number of mRNA transcripts is often normalized relative to the number of cells/transcripts/etc [Wagner et al., 2012], and it is not obvious how well different normalization measures match the distributional assumptions of phylogenetic models of trait evolution. And indeed, there is some empirical evidence to suspect that the assumptions of the independent contrasts method used by Dunn et al. [2018] in their reanalysis of pairwise comparisons were themselves problematic [Begum and Robinson-Rechavi, 2021].

In a recent study, Chen et al. [2019] evaluated the fit of a set of alternative models to gene expression data. This set of models included Brownian motion (BM) [Felsenstein, 1973] and varieties of the Ornstein-Uhlenbeck process (OU) [Hansen, 1997]. Under BM, a phenotypic trait *z* with population mean 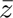 is expected to change over time period *t* according to a random walk such that 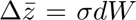, where *dW* is a stochastic process drawn from a normal distribution with variance *t* and mean of 0, which is scaled by the parameter *σ*, such that *σ*^2^ is defined as the evolutionary rate of the BM process. Over time, the variance between replicate lineages (i.e., lineages that share a common ancestor and subsequently had independent evolutionary trajectories) of the phenotypic trait is expected to increase linearly at a rate equal to *σ*^2^. The covariance between replicate lineages is proportional to the amount of shared evolutionary history. The OU process is an extension of the BM model where the mean change in phenotype over some period *t* is described by 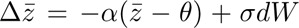, where *α* is some pressure parameter keeping the trait value towards some optimal trait value *θ* with the same random walk *σdW* from BM contributing stochastic divergence. Chen et al. [2019] assessed the utility of phylogenetic models by comparing the relative fit of an alternative set of models using AIC [Akaike, 1974].

However, model selection (using AIC, for instance) is designed to find the model that most closely approximates the generating model [Burnham and Anderson, 2004] balancing accuracy with the additional prediction error that comes with adding free parameters. The fact that a model is favored by model selection does not indicate whether it, or any of the compared models it is compared to, performs well (i.e., is “adequate”), in the sense that the distributional assumptions of the fitted model is consistent with the actual data. This is critical because even the best of a set of models may not adequately describe the structure of variation in the data and conclusions based on an inadequate model may not be reliable. Absolute model performance is typically assessed (when it is) with either parametric bootstrapping [Efron and Tibshirani, 1993] when model parameters are estimated using maximum likelihood, or posterior predictive simulations [Gelman et al., 1996, Rubin, 1984] when parameters are estimated using Bayesian inference. Both parametric bootstrapping and posterior predictive simulations involve simulating new datasets given the model and fitted parameter values and assessing whether the observed data resembles the simulated datasets. If it does, then the model is considered to perform well for the observed dataset (for an overview of methods for assessing the performance of models in the context of evolutionary biology, see Brown and Thomson [2018]).

Pennell et al. [2015] developed an approach, which they implemented in the R package ‘Arbutus’, designed to perform parametric bootstrapping or posterior predictive simulations for phylogenetic models of continuous trait evolution. In brief, the procedure is as follows. First, a model of trait evolution is fit to a comparative dataset using either maximum likelihood, Bayesian inference, or alternatives. Second, the parameter estimates are used to re-scale the branch lengths of the tree such that, if there is a perfect correspondence between the generating and fitted model, whatever the distribution of the data on the real tree, the distribution of the data on the re-scaled “unit tree” will match the expectations of a BM model where *σ* = 1. This re-scaling procedure is valid for any model that satisfies the 3-point condition of Tung Ho and Ané [2014], which includes most models of quantitative trait evolution that have been developed (and all those included in the present study). Since the phylogenetic distribution of gene expression counts is expected to resemble that of a BM model, the phylogenetic independent contrasts (PICs; [Felsenstein, 1985]) computed on the tree would be i.i.d. and *∼ N* (0, 1). Third, various summary statistics are used to compare the distribution of the PICs computed from the real data on the unit tree with the idealized distribution. If the observed summary statistic falls in either tail of the distribution of simulated summary statistics (e.g., *P <* 0.05), the model can be considered to perform poorly (or to be inadequate), because it indicates a substantial mismatch between the distributional expectations of a given model and that of the data. We note that, even if multiple test statistics are used, there no need to perform Bonferroni, False Discovery Rate, etc. correction as there is only hypothesis being tested per gene (*H*_0_ : the data was generated by the fitted model under the estimated parameters). (If one were trying to identify *specific genes* that deviated from the expectations of, say an OU process, this would likely require some type of correction for multiple comparisons.)

Each of these summary statistics measures deviations in the expected distribution of contrasts in unique ways [Pennell et al., 2015]. The statistic *c.var* is the coefficient of variation of the absolute value of the PICs and is a measure of how well a model accounts for rate heterogeneity across a phylogeny. The statistic *d.cdf* is the D statistic from the Kolmolgorov-Smirnov test and measures deviations from the assumptions of normality for the contrasts such as in the case of rapid bursts of phenotypic character change. *S.asr* is the slope of a linear model between the absolute value of the contrasts and the inferred ancestral state of the nearest node to detect if magnitude of a trait is related to its evolutionary rate. The statistic *s.hgt* is the “node height test”, which has been previously used to detect early bursts of phenotypic trait evolution such as in the case of an adaptive radiation [Freckleton and Harvey, 2006, Slater and Pennell, 2014]. The statistic *s.var* is the slope of a linear regression between the absolute value of the contrasts against the expected variances of said contrasts and can be used to detect if the phylogenetic tree used in the fitted model has errors in the branch lengths.

In this paper, we use the approach of Pennell et al. [2015] to assess the performance of commonly used phylogenetic models of evolution for gene expression. We used datasets from previously published studies that leveraged phylogenetic models across a variety of tissues, genes, and species. We have two aims in this paper. First, assessing the performance of phylogenetic models will provide key insights into the general dynamics of gene expression evolution: a number of studies have found that OU models are favored in comparison to BM models — is this because OU processes actually capture the key elements of gene expression evolutionary dynamics, or is it simply because OU models are the best of a poor set of descriptors? Second, we sought to illustrate how techniques for assessing model performance can be applied to individual comparative gene expression studies. Many statisticians [e.g., Gelman et al., 1995] advocate that model-checking should be a routine component of any data analysis yet to our knowledge, this has not been done in any phylogenetic study of gene expression. While in principle, we could use our approach to assess whether the models used to make particular inferences for the particular datasets we used for our analysis (Table 1), doing so would require diving deep into the specific biological question being asked in each (testing some hypotheses will be lean less heavily on the match between a dataset and the assumptions of a model than others) and replicating the exact procedure used in each study — both of these things, while valuable, are beyond the scope of the present work.

**Table 1:**
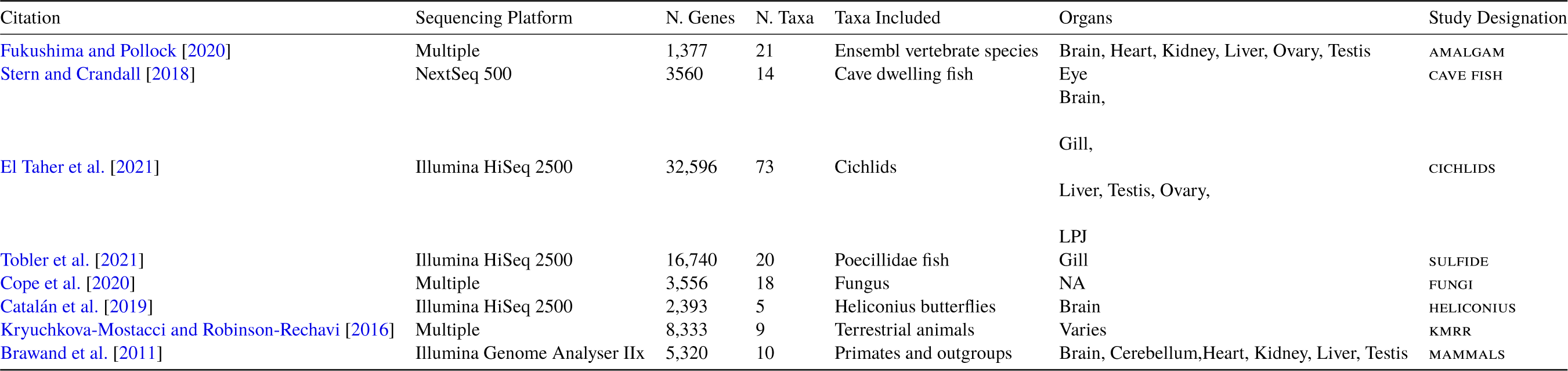
Datasets included in this analysis. Data has to be making use of one of the evolutionary models, provide a phylogenetic tree, and have readily available gene expression data to be used in this analysis.

## Results

### OU models are the best supported model for the majority of genes but adequacy is mixed

In our analyses, we fit three core models: the aforementioned BM, OU, as well as Early Burst (EB) [Blomberg et al., 2003, Harmon et al., 2010]. EB has not, to our knowledge, been applied to gene expression data but we included it because it makes a different set of distributional assumptions, such that it is a useful point of comparison. The EB process, often thought to characterize adaptive radiations [Harmon et al., 2010], is essentially the opposite of an OU model [Uyeda et al., 2015]; the OU model leads to changes to the phenotypic variance being concentrated at the tips of the phylogeny whereas EB concentrates the variance near the root. Mathematically, the EB model is described by an exponential decrease in the rate of evolution through time *t* where some trait mean *z* is determined by Δ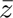(*t*) = *σ*(*t*)*dW*, such that the evolutionary rate at time *t*, 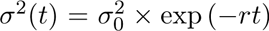, where 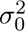 is evolutionary rate at the beginning of the process and *r* describes the decrease in evolutionary rate.

There are two levels of fit we considered for phylogenetic modeling: relative fit — i.e., of the possible models for this set of data, which describes it the best — and absolute fit — i.e., is the model describing the data well? For each of the studies listed in Table 1, we performed a series of analyses that can be summarized along those two tiers. First, we assessed the relative support for each of the three models on each of the genes in the dataset, as measured by AIC weights (following [Harmon et al., 2010]), to determine which of the three models best describes the evolution of that gene’s expression (Figure 1) (See Methods for details). The best fit model for a gene was determined to be the model that minimized AIC. Second, we used Arbutus to measure the performance of the best-fit model for that gene’s data (Figure 1). If multiple tissue types were included, model fit and performance was determined for each tissue type. Since we were both interested in the general distributional properties of mRNA counts across a phylogeny (i.e., we were not looking to explain the evolutionary dynamics of any particular gene) and also wanted to broadly mirror standard practices in the field, we fit models to each gene/tissue combination independently. This assumption, while not correct, allowed us to ask our key questions in a manageable way.

**Figure 1:**
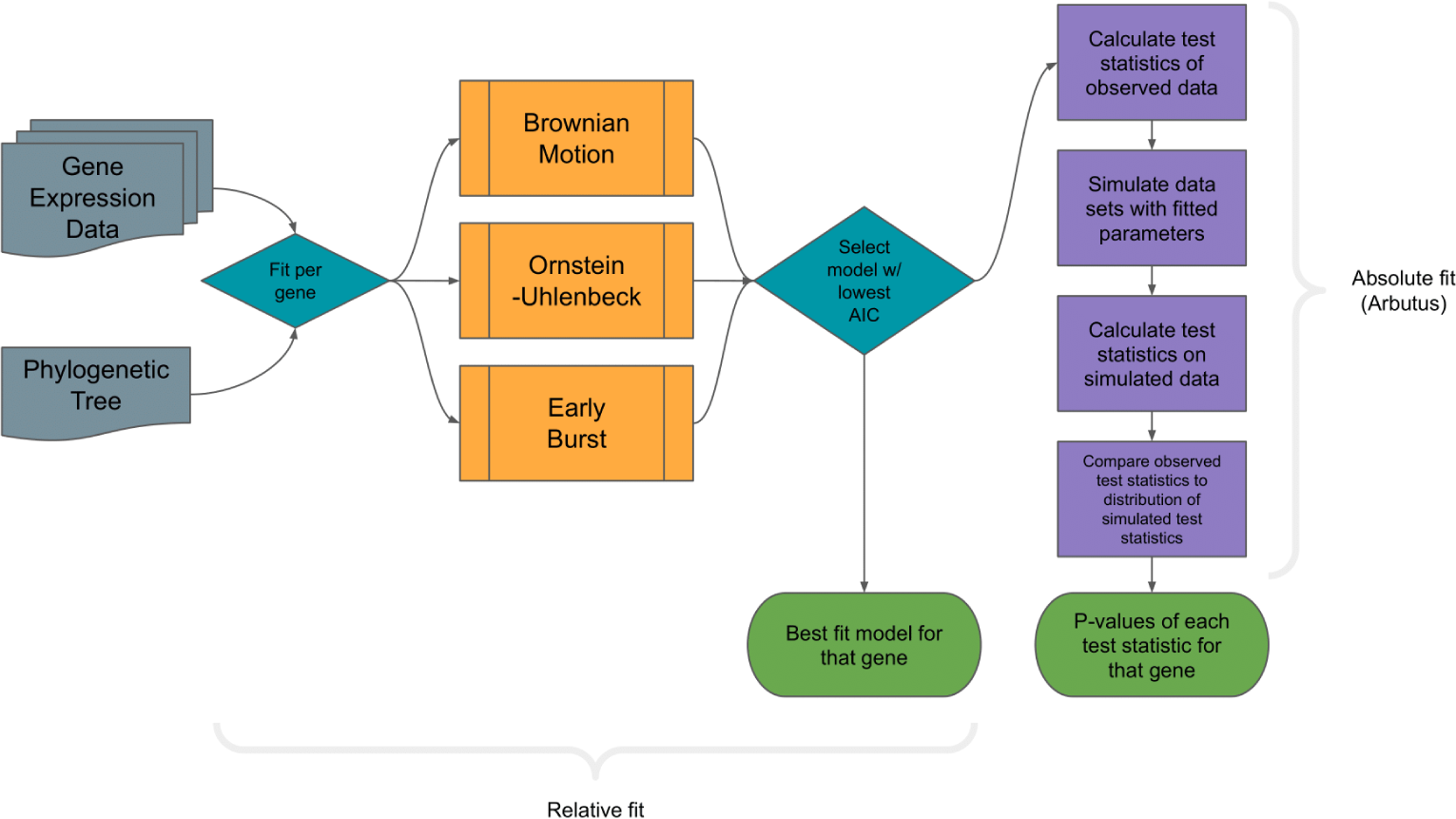
Workflow for determining relative and absolute fit of phylogenetic character models for gene expression data. Data for each gene in a data set is analyzed by first fitting tested PCMs and then testing the best fit model for model adequacy using Arbutus. For data sets with available local gene trees, each gene is paired with its corresponding phylogenetic relationship.

We found that the OU model was the best fit model for 66% (96,307/145,927) of gene/tissue combinations, with noticeable exceptions in heliconius and sulfide where the BM model was the best fit model for 66.5% and 63.8% of genes respectively (Figure 2, Table S1). Notably, the heliconius phylogeny is the smallest included in this study (Table 1) and we have low power to support more complex models. This core result — that OU is better favored over BM — is consistent with several other studies [e.g., Bedford and Hartl, 2009, Chen et al., 2019, Nourmohammad et al., 2017]. Here we are able to go further and ask whether OU is actually describing the data well. We find the data to be mixed: in 61% of cases, the best fit model was found to perform well, as measured by our five summary statistics; for 41.5% of the gene/tissue combinations (60,521/145,927), OU was both the preferred model and a reasonable descriptor of the data. Where the model did perform poorly, it needed to be picked up (i.e., an overabundance of *P* -values *<* 0.05) using the summary statistics *c.var*, *s.asr*, or both (Figure 2, Table S2); this suggests that in cases where models performed poorly, they did so because there was unmodeled heterogeneity in rates of evolution across lineages [Pennell et al., 2015]. So in summary, while there is wide support OU models over alternatives, particularly for the clades with more taxa (where we have more power to detect more complex dynamics), our results reveal that this support is due to a mixture of some genes actually having dynamics that are well-approximated OU processes and some genes having more complicated dynamics for which OU models are only the best of a set of a bad models.

**Figure 2:**
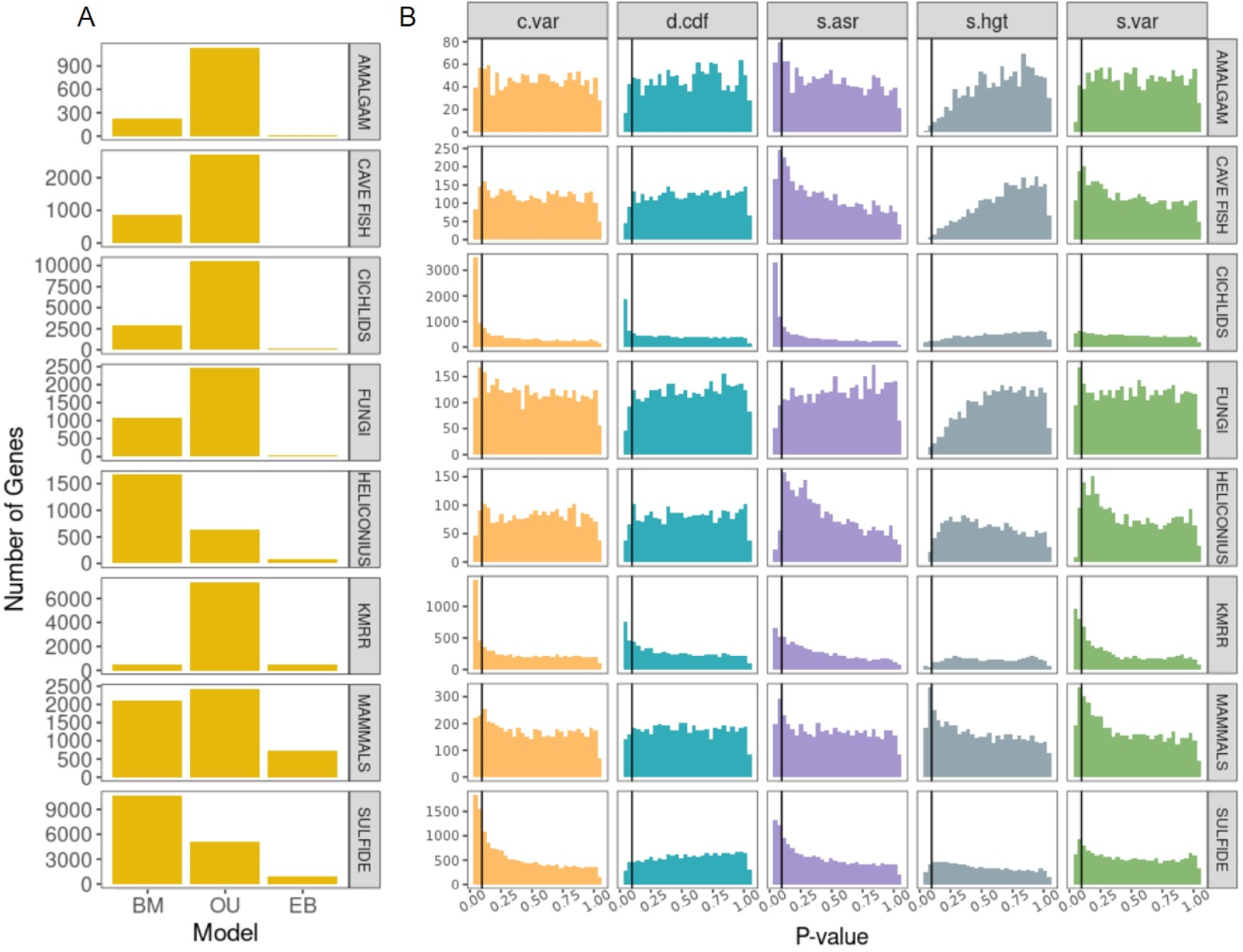
Relative (A) and absolute (B) fit of evolutionary models to the 9 gene expression data sets. Note that the absolute fit was only evaluated on the (relative) best fit model for each gene. Vertical black lines represent the significance cutoff of 0.05, with an expectation of 5% of genes being inadequate by chance. 66% of genes conform to the OU process. In terms of absolute performance, for 61% of genes the best fit model was adequate across all five test statistics. Model failures were primarily prevalent in *c.var* and *s.asr*.

### Normalization does not have much qualitative effect on model performance

All of the datasets we used were from bulk tissue mRNA expression experiments and as such, an appropriate normalization of such counts is critical [Freedman et al., 2021, Wagner et al., 2012, Zwiener et al., 2014] [but see Church et al., 2022, for an alternative approach that does not require normalization]. There was variation in how the counts were normalized in the studies from which they were derived; some being normalized as RPKM (Reads Per Kilobase of transcript, per Million mapped reads) while others as TPM (Transcripts Per Million). For most of the datasets, we did not have access to the original RNAseq counts and as such we could not explore the effects of various transformations on model performance; however, this data was available for the cavefish dataset so we were able to examine this. For this dataset, we looked at three transformations: RPKM, TPM, and CPM (counts per million). In macroevolutionary studies, it is standard practice to log-transform continuous variables, both because this typically means the data better matches the Gaussian assumptions of phylogenetic models and because we are primarily interested in describing phenotypic change on a geometric scale [Houle et al., 2011]. And the same holds true for gene expression data [Diaz et al., 2023] and therefore we only considered log-transformations of RPKM, TPM, and CPM. We find the results to be broadly similar between the various normalization approaches (Fig. S3, Tables S3, S4), which is reassuring. While there may be benefits of alternative normalization schemes in other contexts [Freedman et al., 2021, Zwiener et al., 2014], the patterns we are looking at — interspecific divergences in bulk RNAseq experiments — are so coarse that these differences do not matter much.

### Models fit to species tree have better performance

Models fit to the kmrr data set showed poor performance across the board (Figure 2, Table S2). One major difference between this study and data sets where the models also performed poorly (i.e., fungi, helico-nius, and amalgam), is the type of phylogenetic tree used. Unlike the other studies which each provided the species phylogeny they used for analysis, Kryuchkova-Mostacci and Robinson-Rechavi [2016] instead provided and used gene family phylogenies for each of the genes studied. Comparative analyses of “conventional” phenotypic traits, such as morphology, are typically conducted by using the species tree. However, if the genes underlying the phenotype are in regions of the genome that have different evolutionary histories than the species tree, estimates of phenotypic evolution may be biased. This is true of highly polygenic traits [Hibbins et al., 2023, Mendes et al., 2018] but appears especially problematic for traits that are underlain by a few genes [Hahn and Nakhleh, 2016]. So if the evolutionary models we used actually described evolution quite well, we would expect to see better model performance when using gene trees constructed from the regions of the genome that determine the expression of a particular gene. On the other hand, phylogenetic error, particularly in the branch length estimation, may be particularly acute when estimating trees from small regions, which may introduce an additional set of problems. Unfortunately, we do not know the loci responsible for variation in gene expression for most of the genes so a reasonable approximation would be to use the gene tree of the expressed gene itself as this should be closely linked to the promoter region, whose evolution will likely be important for the evolution of gene expression [Haberle and Stark, 2018, Vaishnav et al., 2022].

Investigating this question comprehensively is beyond the scope of this paper as we do not have access to the original genomic sequence data for all of our datasets. But we did explore this by examining the fungi dataset using both the species tree and local gene trees for all the sampled genes (see Methods for how these trees were constructed). Substituting gene family phylogenies for the species phylogeny reduced the model performance as measured for all test statistics except for *D.cdf*. Two test statistics of note here would be *S.var* and *S.hgt*. The *S.var* statistic will indicate a model is inadequate when there are issues in branch length for the phylogeny used. The number of NA values for *S.hgt* was much higher when using species phylogenies, which could indicate low phylogenetic signal (see [Münkemüller et al., 2012] for discussion of the measurement and interpretation of phylogenetic signal) when using this type of phylogenetic tree (Figure 3). This was confirmed to be the case with Blomberg’s K [Blomberg et al., 2003], where it shows lower K values for genes with NA values in *S.hgt* and thus, lower phylogenetic signal (Figure S2). This higher incidence of NA values arises from the model fitting process. The summary statistic *S.hgt* is the slope of the relationship between the size of the contrasts and the height at which they occur. If an OU model is fit and the *α* parameter is very large, this essentially means that there is no phylogenetic signal. And in this case, the branch lengths leading to the tips of this transformed “unit tree” (see [Pennell et al., 2015] for full mathematical details), will be very long. This means that there will be very little variance in node heights on the transformed trees and it will be therefore impossible to robustly estimate a slope; as such these cases are reported as NAs and excluded from subsequent analysis.

**Figure 3:**
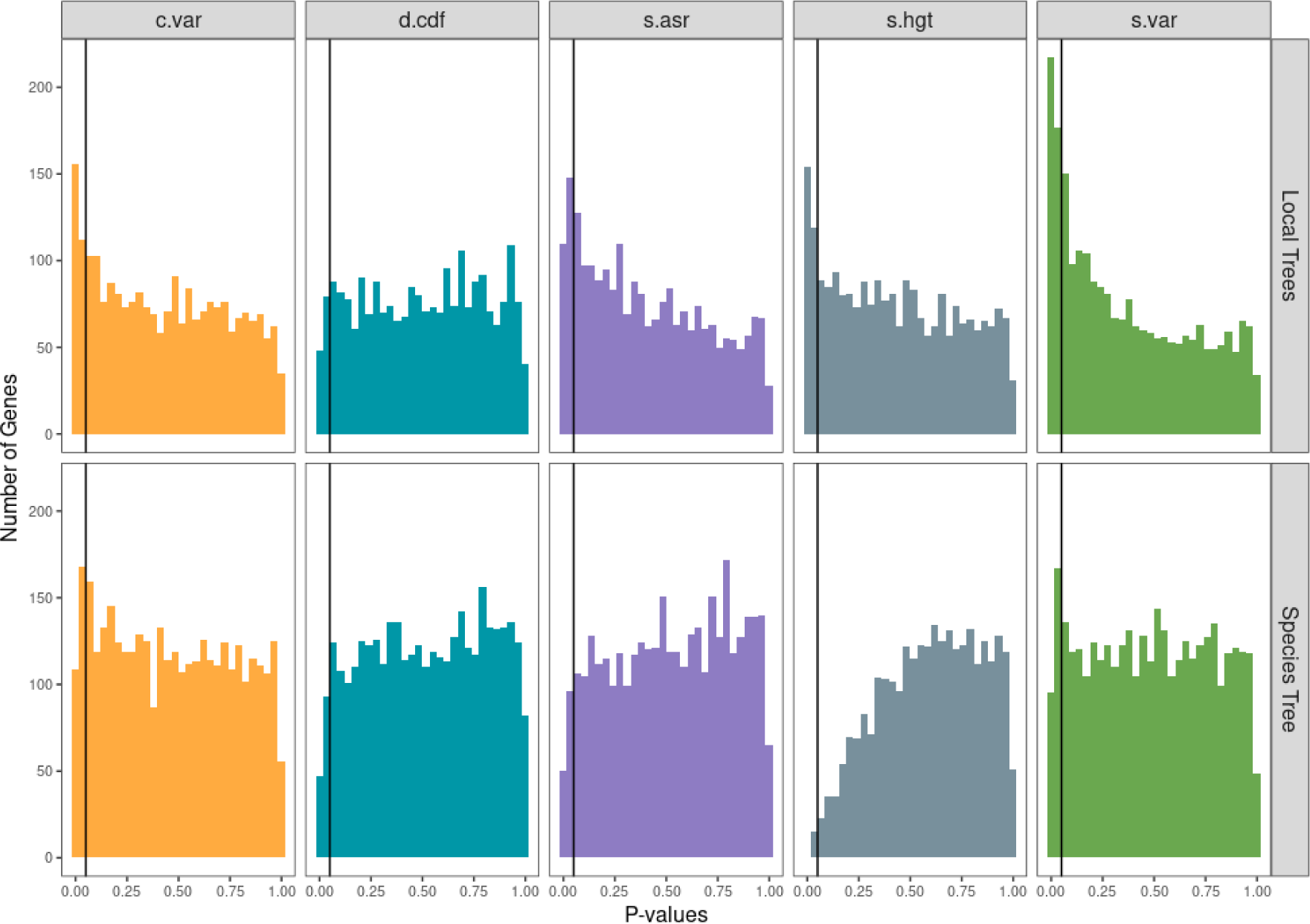
Analysis of generated local gene phylogenies from the fungi data set. Test statistic *P* -values for best-fit models fit with the local gene phylogenies against those fit with the species phylogeny. Models showed poorer performance when they were fit to the gene trees versus the species trees, as measured by the summary statistics *c.var*, *s.asr*, *s.hgt*, and *s.var*.

Taken all together, it seems that for many genes, gene expression has higher phylogenetic signal when models are fit to the local gene trees but overall, models have better performance when they are fit to the species tree, primarily owing to the local trees have a higher frequency of violations detected by the *S.var* summary statistic, which we expect to be violated when there is a lot of branch length error [Pennell et al., 2015].

## Discussion

The relative support of OU models apparent in gene expression datasets — which we also document across 145,927 gene and tissue combinations — is often taken as evidence for the role of stabilizing selection [Bedford and Hartl, 2009, Chen et al., 2019, Nourmohammad et al., 2017] but there are reasons to be critical about this interpretation [Price et al., 2022].

First, there are technical challenges; these include difficulties in estimating the parameters of an OU process and conducting model comparisons [Cooper et al., 2016] (but see [Grabowski et al., 2023]) as well as various experimental artifacts, which together can potentially create a lot of measurement error [Price et al., 2022]. This is particularly problematic for comparing the fits of evolutionary models because unmodeled measurement error will tend to data that may appear more OU-like [Cooper et al., 2016, Pennell et al., 2015, Silvestro et al., 2015]. Ideally, this can be dealt with by jointly modeling macroevolutionary dynamics and the processes that generate biological error within species [Rohlfs et al., 2014, Rohlfs and Nielsen, 2015] but this requires multiple measurements per gene per lineage and this was not available for most of the datasets we analyzed. The best we could do was to estimate a standard error for the estimates of the mean expression (see Methods for details) and include this as a fixed parameter when we fit the phylogenetic models. We emphasize that these challenges apply equally to the types of morphological traits (e.g., body size) that phylogenetic comparative methods are usually applied to. And indeed, it is notable that our results reveal that despite the likely differences in genetic architecture between morphological phenotypes and gene expression levels, their general phylogenetic distribution is broadly similar (i.e., the general findings here closely match that of Pennell et al. [2015]).

Gene expression is different from morphological data in that theory suggests that we do not necessary expect gene expression to be under stabilizing selection. In all of the studies we re-analyzed, gene expression was treated more-or-less synonymously with mRNA abundance. However, the protein abundance is much closer to the phenotype. The mRNA abundance of a cell is, in many studies (and often implicitly) treated as a proxy for protein abundance. Many studies have compared mRNA and protein abundances across genes (and in a few cases, species) and found varying level of correlation [Ba et al., 2022, Becker et al., 2018, Gygi et al., 1999, Khan et al., 2013, Laurent et al., 2010, Marguerat et al., 2012, Wang et al., 2019]. Studies have also found protein abundances to be more phylogenetically conserved than gene mRNA abundance. These observations have been attributed to the presence of compensatory evolution, in which selection for changes to the protein abundances can lead to selection for changes in the transcription rate but which can also be compensated for by changes to the translation or degradation rate of the gene products [Khan et al., 2013, Laurent et al., 2010, Schrimpf et al., 2009, Wang et al., 2020]. This verbal hypothesis was recently formalized in a theoretical study by Jiang et al. [2023]; they found that when there is stabilizing selection on the protein abundance, such compensatory evolution was indeed expected and this would typically result in evolution of mRNA that resembled patterns produced by a BM model albeit with lower rates of divergence than expected under pure genetic drift (see [Jiang and Zhang, 2020] for more on this point). (Morphological phenotypes, in contrast, are more likely to be a direct target of selection, stabilizing or otherwise.)

In light of all of this, our finding that simple models, and OU models in particular, are often (i.e., in the majority of cases) reasonable descriptors of the evolutionary dynamics of gene expression can be interpreted in multiple ways. On one hand, this could suggest that inferences are being driven by unaccounted for measurement error. And if this is the case, we should be suspect of many previous empirical claims that are dependent on inferences from these models. On the other hand, if one believes that technical artifacts are reasonably well accounted for (we included an estimate of measurement error in our study and also demonstrated that the normalization scheme employed did not qualitatively affect our results), this suggests that perhaps there is stronger stabilizing selection on mRNA levels (i.e., to match the selective optima of the corresponding protein levels) than verbal and quantitative models (see Jiang et al. [2023] and references therein) of gene expression evolution predicted.

For the genes where the simple models performed poorly, this was typically because they failed to capture important sources of variation across lineages. Unmodeled heterogeneity in the process may confound inferences. For example, if one is testing whether gene expression levels for a particular gene remain near some macroevolutionary optima or optimum (*sensu* [Arnold et al., 2001]) and include only a single optima (when the data implies there are multiple), the inferred macroevolutionary stability (within an evolutionary regime) will be greatly underestimated. In this paper, we only considered the adequacy of relatively simple models of gene expression evolution. This had the advantages of allowing us to characterize the general features of a wide variety of datasets in a coherent way and because it served as a straightforward illustration of how researchers could incorporate tests of model performance into the comparative gene expression workflow. However, it is likely that, for many genes, there will be variation across the phylogeny in the optimal level of gene expression and in the rates at which gene expression evolves. And accordingly, a number of recent studies (including some of original publications from which our datasets were derived) used multi-rate or multi-optimum variants of the OU process (e.g., [Brawand et al., 2011, Catalán et al., 2019, Chen et al., 2019, Fukushima and Pollock, 2020, Stern and Crandall, 2018, Tobler et al., 2021]). (Indeed, Grabowski et al. [2023] argue that this is the primary use-case for OU models.) This is important as a previous analysis by Chira and Thomas [2016] of morphological phenotypes found that when the generating process was a multi-rate evolutionary model, fitting single-rate models to the data (as we have done here) would lead to poor model performance, as detected by the same summary statistics with which violations were commonly detected in our data – and that including multi-rate processes often led to better relative fit compared to single rate models and better model performance on absolute terms. All of these models satisfy the 3-point condition of Tung Ho and Ané [2014] and thus the approach of Pennell et al. [2015] illustrated here, could be applied to check the performance of models in such studies. We did not do this here because we would need to decide on an *a priori* method for assigning branches of the tree to different evolutionary regimes [Beaulieu et al., 2012] or estimating the regimes from the data [Uyeda and Harmon, 2014] but the most relevant approach would really depend on the details of the specific hypotheses being tested — and again, re-evaluating the claims of past studies was beyond the scope of the present work.

An additional factor that will likely affect model performance is the size of the dataset (in terms of numbers of taxa). As gene expression data is still relatively expensive to collect (i.e., compared to many morphological traits), the size of many phylogenetic comparative studies of gene expression is relatively modest by modern macroevolutionary standards. As more and more taxa are included, the greater the chances that there will be substantial heterogeneity in the evolutionary process. It will also be the case that as datasets get larger, there will be more evidence to detect deviations from the assumptions of a model. Unfortunately, when assessing model performance it is rather difficult to disentangle these two factors and whether the distinction matters or not will depend on the research question [Pennell et al., 2015]. Thus, we suspect that the reasonably good performance of relatively simple models may be due, at least in part, to the modestly sized datasets that we analyzed.

And expanding out from the particular analyses and results presented here, we hope to encourage researchers to include the evaluation of model performance as part of their comparative gene expression studies. The approach of Pennell et al. [2015] is general in that it can be adapted to evaluate performance for a wide array of phylogenetic models for continuous traits. Assessing the absolute performance of a phylogenetic model in a comparative gene expression study can provide more confidence in the results of the analyses — if the models used broadly perform well — or suggest new models that should be considered — if they do not. We, like many others, are excited that comparative gene expression studies are increasingly being conducted in a phylogenetic context. There are many important evolutionary questions that we may finally be able to address with this data [Price et al., 2022, Smith et al., 2020]. We hope this contributions aids in this work but helping researchers ensure that their inferences regarding these questions are on a sure footing.

## Methods

### Data

We aimed to explore model performance across a variety of different studies, including a range of taxa, tissues, and genes. To focus on relevant studies, we prioritized studies according to two criteria: first, that the originating study made use of at least one of the evolutionary models being assessed in this analysis and second, where the gene expression data and phylogenetic tree used in the study were readily available. The studies gathered in this process range in both number of genes and species analyzed as well as taxa included and tissues sampled (Table 1).

### Analysis of Model Fit

We log-transformed normalized gene expression values for all data sets (see below for details on normalization) before we evaluated model fit and adequacy to facilitate cross-species comparisons. For every gene-tissue combination, we used the `fitContinuous()` function in geiger [Pennell et al., 2014] to fit BM, OU, and EB to the comparative gene expression dataset. When a species tip was missing data for a gene, that tip was excised before performing fitting and adequacy measurement. If data sets included multiple samples per species, the mean expression was used and an error term equivalent to the standard error of the gene expression data for that species was used for that gene. Relative model fit was assessed on a per-gene basis, with each gene being assigned one model with the best fit; i.e., the model with the lowest AIC score as calculated by the model-fitting process. We then plotted best-fit models using the ggplot R package [Wickham and Wickham, 2016]. Model adequacy was calculated using best-fit model parameters calculated in the previous step using the Arbutus R package [Pennell et al., 2015].

### Evaluating the effect of standardization

To measure the effect of normalization type on model adequacy, we compared model adequacy between RPKM, TPM, and CPM values for the cave fish data set (Table 1). We quality trimmed these reads using Trimmomatic [Bolger et al., 2014] and performed alignment and quantification using the Trinity pipeline [Grabherr et al., 2011], producing both RPKM, TPM, and CPM values for all genes included. To compare adequacy we then performed adequacy analysis as explained above for all normalization methods.

### Local Gene Phylogeny Construction

We generated gene family phylogenies for the Cope et al. [2020] data set using protein sequences downloaded from the Ensembl database [Howe et al., 2021] via the biomaRt package [Durinck et al., 2022]. We then aligned downloaded sequences using MAFFT [Katoh and Standley, 2013] and assembled them into phylogenetic trees using FastTree [Price et al., 2010], which uses a minimum evolution model to build trees. We fit chronograms to gene trees to make them ultrametric using penalized likelihood as implemented in the ape package with the chronos function [Paradis and Schliep, 2019]. We implemented this in a single Snakemake [Mölder et al., 2021] pipeline.

### Testing for Phylogenetic Signal

Phylogenetic signal was compared for genes with NA values in the S.hgt metric against genes with real numerical values using the phytools R package [Revell, 2012]. Results were plotted for both the K-statistic [Blomberg et al., 2003].

## Data and Code Availability

All R scripts, pipelines, and data used in this analysis can be found in or redirected from the following GitHub repository: https://github.com/fieldima/adequacy_of_PCMs.

## Acknowledgements

We thank Paul Pavlidis, Keith Adams, Alex Cope, members of the Pennell and Pavlidis labs, and three anonymous reviewers for their comments on the work and the manuscript. Casey Dunn and Felipe Zapata provided additional insights into the problems addressed here. M.P. was supported by a NSERC Discovery Grant. Research reported in this publication was supported by the National Institute of General Medical Sciences of the National Institutes of Health under award number R35GM151348.

## Supplementary materials

**Table S1:**
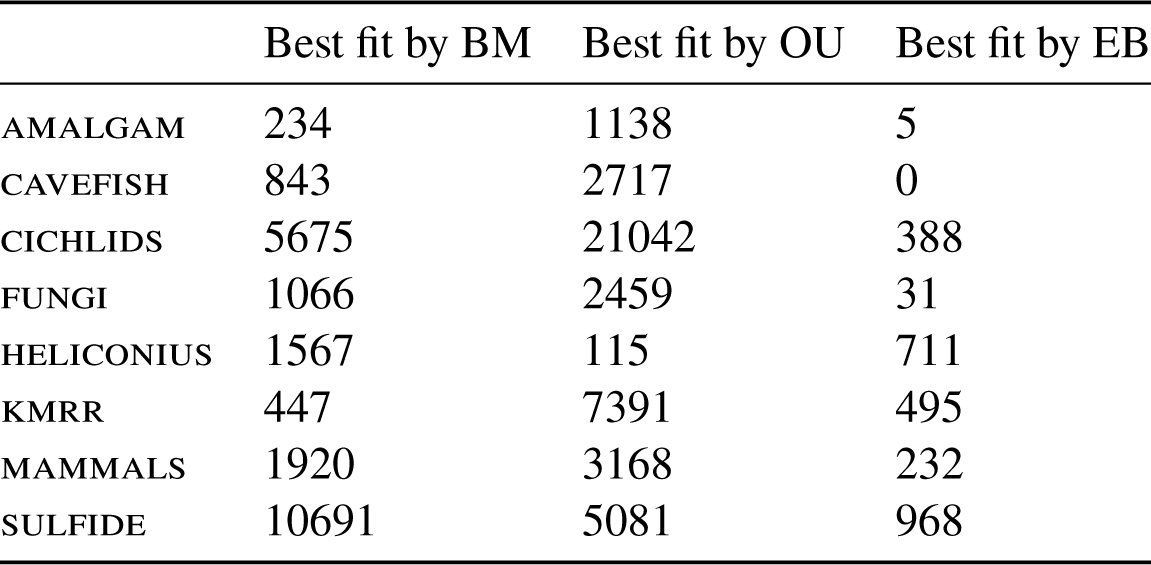
Number of genes in each dataset best fit to three models.

|  | Best fit by BM | Best fit by OU | Best fit by EB |
| --- | --- | --- | --- |
| AMALGAM | 234 | 1138 | 5 |
| CAVEFISH | 843 | 2717 | 0 |
| CICHLIDS | 5675 | 21042 | 388 |
| FUNGI | 1066 | 2459 | 31 |
| HELICONIUS | 1567 | 115 | 711 |
| KMRR | 447 | 7391 | 495 |
| MAMMALS | 1920 | 3168 | 232 |
| SULFIDE | 10691 | 5081 | 968 |

**Table S2:** Percentage of genes in each dataset for which the best-fit model performed poorly (*P <* 0.05) as measured by our five test statistics.

| Study | *c.var* | *d.cdf* | *s.asr* | *s.hgt* | *s.var* |
| --- | --- | --- | --- | --- | --- |
| AMALGAM | 7.2% | 4.5% | 10.1% | 0.6% | 4.5% |
| CAVEFISH | 6.5% | 3.9% | 11.6% | 0.2% | 8.3% |
| FUNGI | 7.8% | 4.0% | 4.1% | 0.6% | 7.4% |
| HELICONIUS | 6.0% | 4.8% | 3.5% | 1.0% | 5.1% |
| KMRR | 23.1% | 13.3% | 15.2% | 1.9% | 14.0% |
| MAMMALS | 17.9% | 7.2% | 15.6% | 1.7% | 3.7% |
| SULFIDE | 20.2% | 4.4% | 15.2% | 6.4% | 9.2% |

**Table S3:**
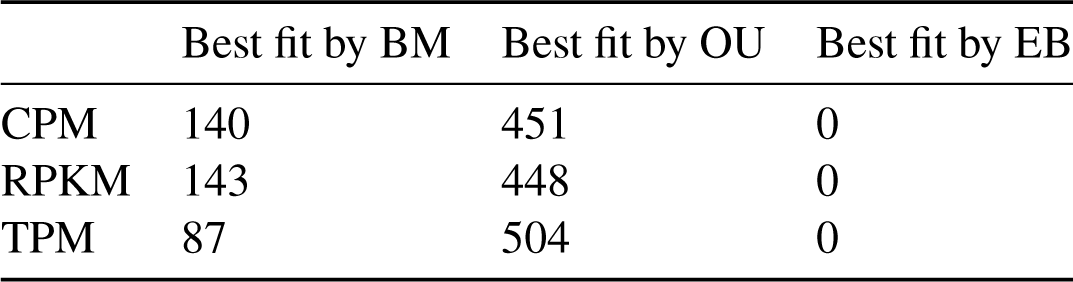
The number of genes in the cavefish dataset best fit to each model when different normalization methods (CPM, RPKM, TPM) are used.

|  | Best fit by BM | Best fit by OU | Best fit by EB |
| --- | --- | --- | --- |
| CPM | 140 | 451 | 0 |
| RPKM | 143 | 448 | 0 |
| TPM | 87 | 504 | 0 |

**Table S4:**
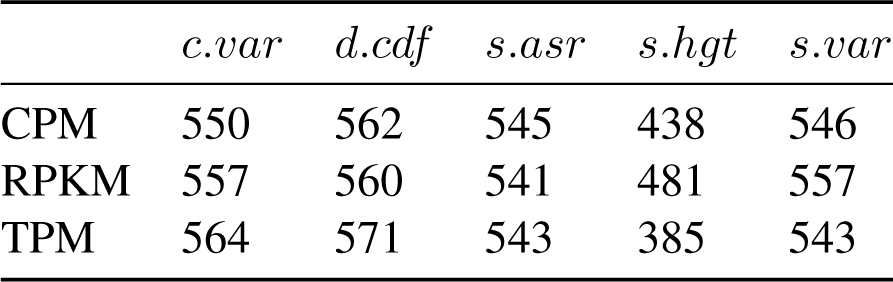
The number of fully adequate genes in the cavefish dataset based on different testing statistics when different normalization methods are used.

**Figure S1:**
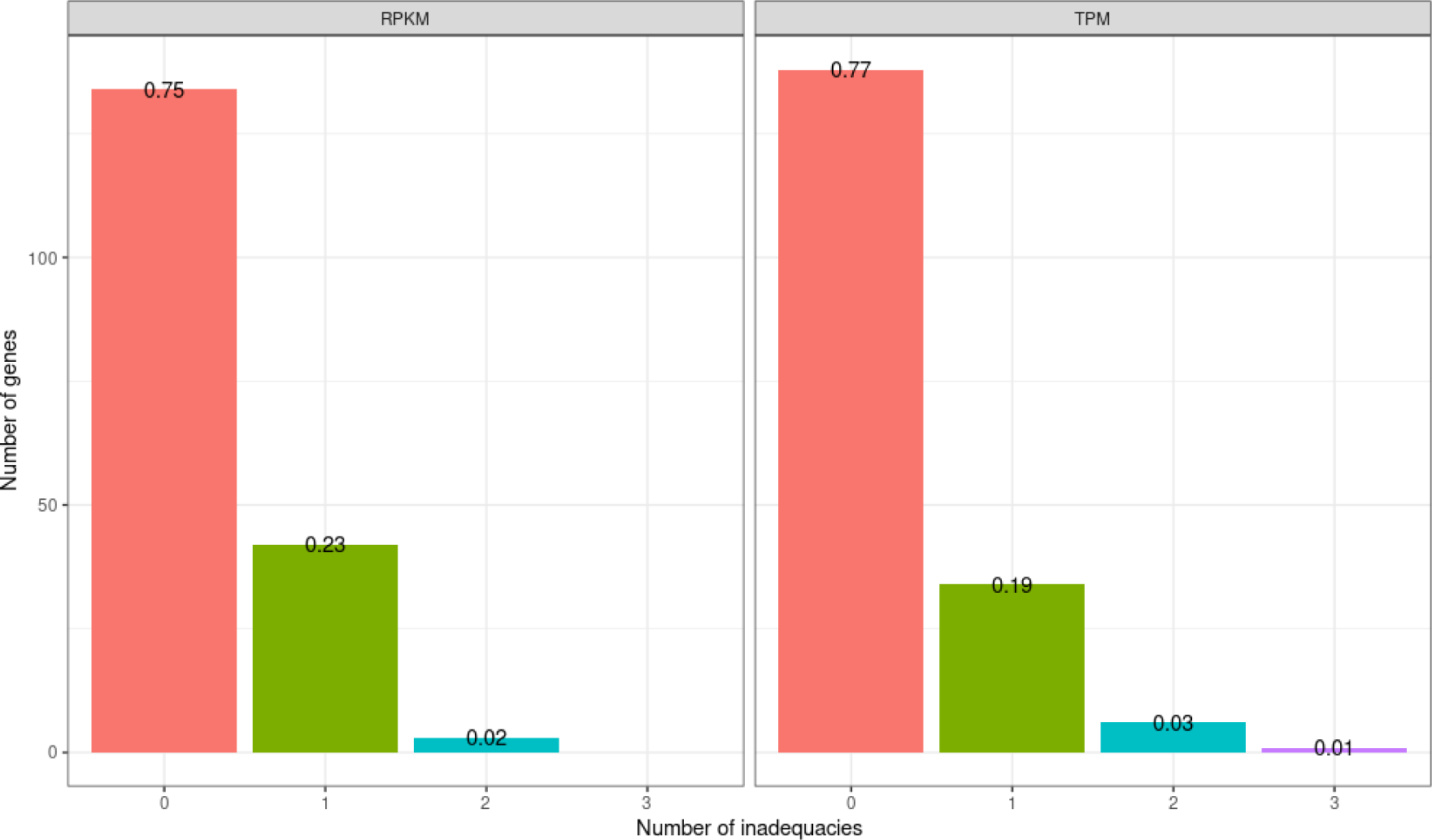
Proportion of inadequate genes for RPKM (left) and TPM (right) normalized reads. Raw reads from the cavefish data set were normalized into RPKM and TPM values and then the best fit model was analyzed for model adequacy via ARBUTUS. The proportion of genes with zero, one, or two inadequacies was nearly identical between both modes of normalization.

**Figure S2:**
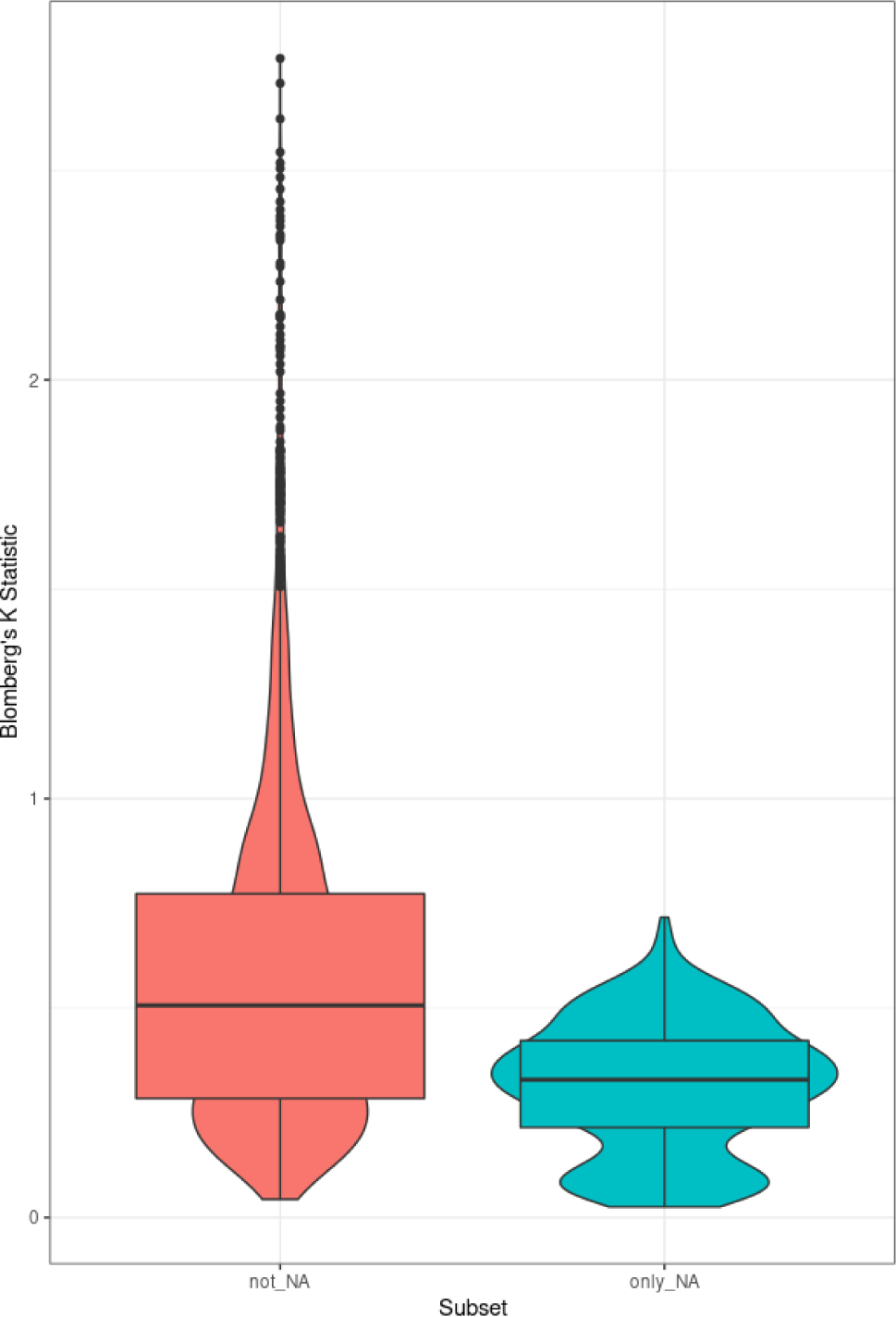
Blomberg’s K statistic for genes with NA values (left) and non-NA values (right) in the S.hgt test statistic. Genes with NA values have a lower K statistic on average than genes without.

**Figure S3:**
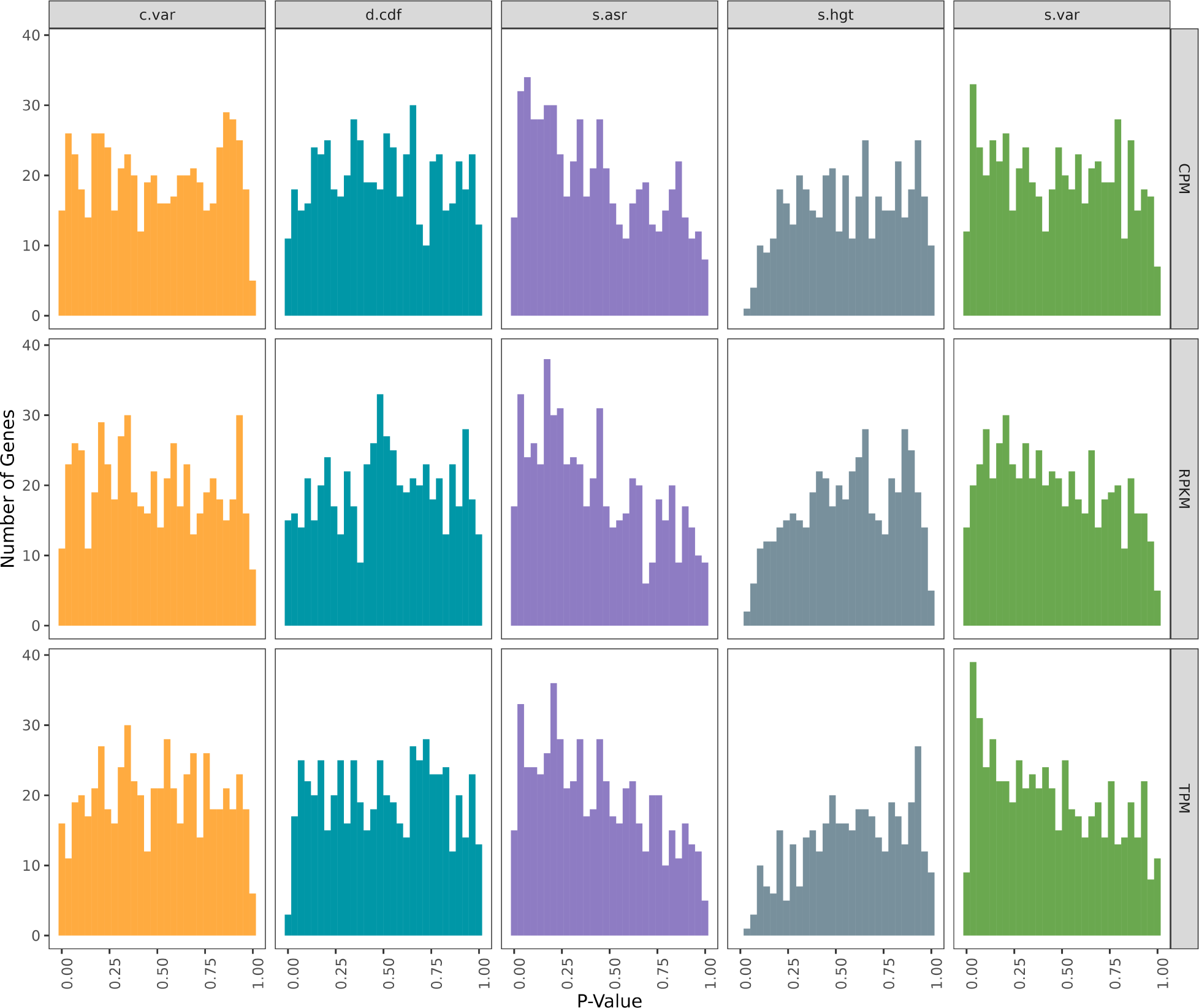
Absolute fit of evolutionary models to the cavefish data set with different normalization methods used. The absolute fit was only evaluated on the (relative) best fit model for each gene. Vertical black lines represent the significance cutoff of 0.05, with an expectation of 5% of genes being inadequate by chance.

## Notes

### Competing Interest Statement

The authors have declared no competing interest.

### Summary of Updates

The introduction and discussion have been restructured and some additional (supplemental) analyses have been added.

